# Trait Convergence Dominates Community Assembly Across Contrasting Himalayan Habitats: Integrating Above- and Belowground Strategies With Phylogenetic Structure

**DOI:** 10.1101/2025.08.12.669809

**Authors:** Jiří Doležal, Ondřej Mudrák

## Abstract

Understanding the mechanisms that structure plant communities is fundamental for predicting biodiversity responses to global change. Functional trait–based approaches can reveal assembly processes by linking species’ adaptations to community-level patterns, with convergence typically signalling strong environmental filtering and divergence reflecting niche partitioning or competitive exclusion. In extreme environments such as cold, arid mountains, it remains unclear whether persistence is driven primarily by abiotic filtering (low temperatures) or by competition for scarce resources (soil moisture, nutrients) that promotes niche differentiation. Disentangling these mechanisms requires an integrated framework combining multidimensional trait data with phylogenetic structure, enabling separation of adaptive responses from evolutionary constraints in shaping community composition. We analysed community assembly across 482 sites in six contrasting Himalayan habitats: salt marshes, wet grasslands, shrublands, steppes, screes, and alpine tundra, spanning 3,700–5,850 m. Using community-weighted means (CWM) and standardized effect sizes (SES) for 13 above- and belowground traits, including leaf morphology, stem anatomy, clonality, root carbohydrates, and nutrient storage, alongside a multi-locus phylogeny of all species, we linked functional and phylogenetic structure with species richness. Most communities showed strong functional convergence (negative SES) in stress-tolerant traits (e.g., high Leaf Dry Matter Content, Root Phosphorus Content, and Root Non-structural Carbohydrates), particularly in species-poor habitats such as salt marshes and alpine tundra. Divergence (positive SES) was more common in species-rich wet grasslands and steppes, where milder conditions likely facilitate niche differentiation. Phylogenetic overdispersion frequently accompanied functional convergence, indicating repeated evolution of similar trait syndromes in distantly related lineages under shared environmental constraints. Assembly processes had contrasting effects across traits: in alpine and scree communities, conservative traits converged while clonality and δ^15^N diverged, suggesting parallel action of filtering and differentiation; in nutrient-enriched systems, convergence in Specific Leaf Area coincided with divergence in root traits, highlighting trait-specific responses to stress and competition. We conclude that environmental filtering dominates Himalayan community assembly but is frequently modulated by niche differentiation. The consistent decoupling between functional and phylogenetic diversity demonstrates that only by integrating multidimensional trait data with phylogeny can we disentangle adaptation from evolutionary constraint and robustly predict biodiversity responses to environmental change.

## 1 Introduction

Unravelling the processes that assemble plant communities and shape their functional and phylogenetic diversity is central to understanding how biodiversity is structured, maintained, and responds to environmental change (Grime, 2006; Garnier et al., 2015; de Bello et al., 2021). Functional traits (morphological, physiological, or phenological characteristics influencing organismal performance) are fundamental to explaining species composition and interactions within communities (Violle et al., 2007; Lavorel & Garnier, 2002; Díaz et al., 2007; Funk et al., 2017). In particular, analyses of trait dispersion can reveal the relative importance of assembly processes such as environmental filtering, niche differentiation, and neutrality (Keddy, 1992; Kraft et al., 2008; Garnier & Navas, 2012). Understanding the interplay between these processes is especially relevant in ecosystems under intense abiotic stress, where drought, nutrient limitation, and temperature extremes act as powerful ecological filters (Doležal et al., 2016). Yet whether plant persistence in such conditions is driven primarily by abiotic filtering or by competition for scarce resources remains largely unresolved (McGill et al., 2006). This uncertainty is compounded by the scarcity of studies that integrate various traits with phylogeny, particularly in the functionally and phylogenetically underexplored ecosystems such as the Himalayas.

Functional trait–based approaches thus offer a powerful means of linking organismal adaptations to community-level structure and dynamics (de Bello et al., 2021). Trait convergence, where co-occurring species share similar strategies, typically indicates strong environmental filtering selecting for stress-tolerant traits, whereas trait divergence may reflect niche partitioning or competitive exclusion, with species adopting distinct strategies to reduce overlap (Mason et al., 2011). These processes often act concurrently or at different spatial scales, with their relative importance shifting along environmental gradients (Weiher & Keddy, 1995; Mayfield & Levine, 2010; de Bello et al., 2021). Disentangling them requires an integrated framework that combines multidimensional trait data with phylogenetic structure, enabling the separation of adaptive responses from evolutionary constraints in shaping community composition.

Mountain systems, and especially the Himalayas, provide exceptional natural laboratories for exploring these mechanisms due to their sharp climatic, topographic, and edaphic gradients (Dvorský et al., 2017; Dolezal et al., 2018). The western Himalaya harbors a mosaic of vegetation types—including salt marshes, wet grasslands, shrublands, dry steppes, mobile screes, and alpine tundra—that span extreme variation in abiotic stress, productivity, and successional stage (Dvorský et al., 2011). Previous research in this region suggests that cold and drought stress act as strong environmental filters promoting conservative ecological strategies such as low stature, high leaf dry matter content (LDMC), and nutrient and water retention (Dvorský et al., 2011; Binter et al., 2025). While these traits tend to converge in stressful environments, phylogenetic overdispersion often accompanies them (Le Bagousse-Pinguet et al., 2018), indicating that similar strategies have evolved independently across diverse lineages (Webb et al., 2002; Craven et al., 2018).

Trait dispersion patterns, therefore, serve as diagnostic tools to assess the balance between filtering and differentiation (de Bello et al., 2021). Stress-dominated or early-successional habitats typically show convergence in traits, while more productive or less constrained habitats may favor divergence due to higher resource availability and interspecific competition (MacArthur & Levins, 1967; Mudrák et al., 2016). Notably, convergence can also arise in productive settings through competitive dominance of functionally similar species, particularly when asymmetric competition limits trait variance (Götzenberger et al., 2012; Mason et al., 2011). Yet, the extent to which these trait patterns are mirrored or masked by phylogenetic structure remains poorly documented for Himalayan vegetation types.

Crucially, different dimensions of the plant phenotype are shaped by different ecological filters. Aboveground traits like plant height, SLA, and leaf nutrient content often reflect growth potential and light competition (Grime, 2006; Garnier et al., 2015), while belowground traits, including root nutrient content, storage capacity, and clonality, reflect persistence, resource acquisition, and tolerance to disturbance (Klimešová et al., 2024; Doležal et al., 2019a). In the Himalayas, acquisitive strategies with tall, fast-growing species often dominate in steppes and shrublands (Doležal et al., 2019b; Chondol et al., 2025), whereas alpine plants rely on conservative syndromes, including compact growth, high LDMC, and carbohydrate reserves for survival under freezing stress (Doležal et al., 2019c; Binter et al., 2025; Chlumská et al., 2023).

Belowground morphology and physiological traits are particularly relevant for plant persistence in cold and drought-prone systems (Klimešová et al., 2011). Storage organs and clonality enable plants to buffer against environmental variability and disturbance, as well as to exploit heterogeneous microsites. For instance, de Bello et al. (2011) showed that in East Ladakh’s alpine zones, clonal growth forms exhibit both convergence due to environmental filtering and divergence driven by niche partitioning. This points to the dual role of clonality in assembly processes, simultaneously enhancing persistence and facilitating coexistence through spatial differentiation (Klimešová & Herben, 2024). Yet, clonality and storage traits have rarely been integrated into broad-scale functional–phylogenetic analyses in Himalayan ecosystems, leaving their macroecological roles largely unresolved.

Phylogenetic data, when integrated with trait analyses, can further reveal whether observed trait patterns result from shared ancestry or convergent adaptation (Hähn et al., 2024). High phylogenetic conservatism in trait expression may indicate evolutionary constraints, while convergence among distantly related taxa implies similar selection pressures shaping functionally analogous strategies (Webb et al., 2002; Cavender-Bares et al., 2009; Craven et al., 2018). In several Himalayan plant communities, functional similarity coincides with phylogenetic divergence (Le Bagousse-Pinguet et al., 2018; Doležal et al., 2019c), suggesting that environmental filtering has repeatedly selected for analogous traits in unrelated lineages (Doležal et al., 2019a; Binter et al., 2025). However, the simultaneous assessment of above- and belowground traits alongside phylogenetic information across multiple Himalayan habitats is still lacking, impeding a comprehensive view of trait coordination and trade-offs.

In this study, we analyzed functional composition and diversity across six distinct plant communities in the western Himalaya using a multidimensional trait dataset including leaf, root, stem anatomical, and whole-plant traits. We quantified community-weighted means (CWMs) and standardized effect sizes (SES) of functional diversity to test two main hypotheses: (1) whether communities differ in average trait composition; and (2) whether trait dispersion patterns indicate convergence (environmental filtering), divergence (niche differentiation), or neutrality. By also incorporating species phylogeny, we evaluated whether functional patterns align with evolutionary relatedness. We expected convergence in stress-prone habitats like salt marshes and alpine tundra, and greater trait divergence in more productive or structurally complex communities such as shrublands and steppes.

## 2 Material and Methods

### 2.1 Study Area and Vegetation Sampling

The study was conducted in the western Himalaya, primarily in eastern Ladakh, across an area of approximately 40,000 km^2^ (32.47-34.46N, 75.50-78.60E). Sampling spanned a steep elevational gradient from 3,700 to 5,850 m a.s.l., capturing a wide range of climatic and edaphic conditions. Along this gradient, mean annual temperature declines sharply from ∼13°C at lower elevations to –13°C at the highest sites while annual precipitation increases from ∼50 mm to ∼250 mm (Dvorský et al., 2015; Macek et al., 2021). Seasonal temperature amplitude averages 46 ± 5°C, and precipitation is mainly in the form of snow at high elevations and sporadic rainfall at lower elevations (Dvorský et al., 2015, 2016).

The sampled vegetation encompassed six ecologically distinct communities (Fig. 1), each representing a unique combination of elevation, substrate, and moisture availability. Salt marshes occur at lower elevations on saline or alkaline soils and are dominated by low-stature, clonal, stress-tolerant species. Wet grasslands are found in flat or gently sloping sites with high soil moisture, often along streams, and support dense herbaceous vegetation. Steppes are open, arid habitats with well-drained, gravelly or sandy soils and sparse drought-adapted shrubs and grasses. Shrublands are located on rocky mid-elevation slopes and are dominated by cushion-forming or hemispherical shrubs. Screes consist of steep, unstable slopes with coarse rock debris and support specialized, mechanically robust plant species. The alpine zone, located at the uppermost elevations, is characterized by extreme cold, high wind exposure, and nutrient limitation, and supports mat-forming and cushion vegetation adapted to short growing seasons.

**Figure 1:**
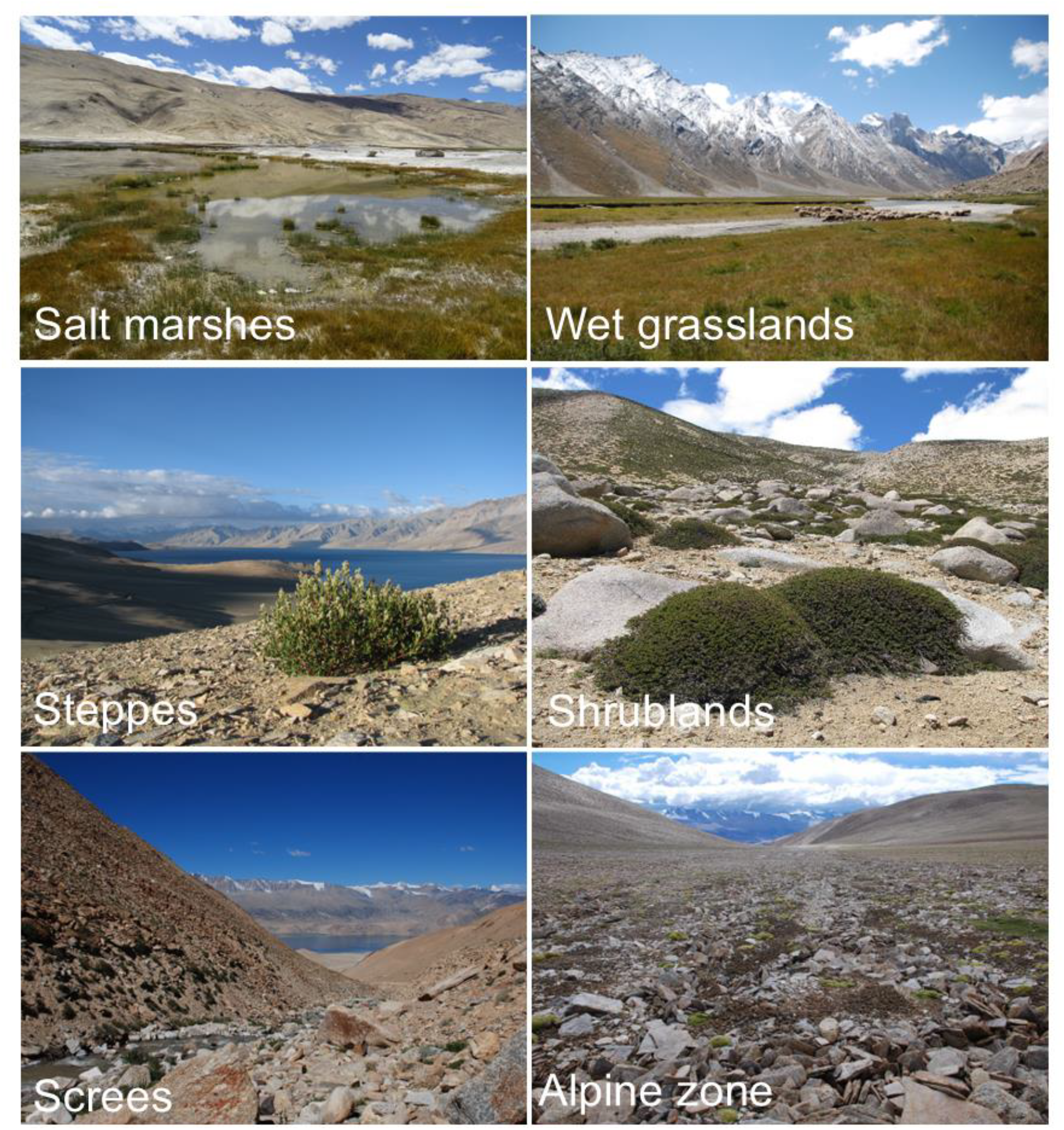
Representative images of the six studied Himalayan habitats: salt marshes (SM), wet grasslands (WE), steppes (ST), shrublands (SH), screes (SC), and alpine zone (AL). These habitats differ in topography, moisture availability, and disturbance, shaping distinct plant communities.

A total of 482 vegetation relevés were recorded across these six communities using standardized phytosociological protocols, encompassing 338 species of vascular plants. Each relevé was established within a 10 m^2^ plot, where all vascular plant species were identified and their cover visually estimated. Total plant cover, species richness, vegetation height, and community structure were also recorded. Community identity was confirmed using dominant species composition and physiognomic characteristics. Environmental data, including elevation, slope, and soil conditions, were documented to assess habitat-specific variation and environmental filtering across the gradient.

### 2.2 Plant trait collection and measurement

To capture functional trait variation across environmental gradients, we sampled over 7,800 individuals representing 338 vascular plant species recorded in 482 vegetation relevés throughout Ladakh (western Himalaya) between 2012 and 2017. Sampling occurred during peak growing seasons (July–August) to ensure trait measurements reflected optimal phenological stages and were minimally affected by drought or senescence. Due to logistical constraints in sampling across a 40,000 km^2^ area and an elevational range of >2,000 m, trait measurements were conducted over multiple years. Most species were long-lived perennials with root and rhizome systems persisting for decades (Chondol et al., 2023), ensuring that measured traits represented long-term ecological strategies rather than transient states.

For each species, we selected 20–30 healthy, undamaged individuals from populations growing near their elevational optima, defined by peak density and abundance, to best reflect species’ trait syndromes under locally adapted conditions. Each plant was excavated with intact root systems, measured for height, and separated into leaves, stems, reproductive organs, and belowground parts (roots or rhizomes). After cleaning, we immediately fixed 3–5 cm segments of roots/rhizomes in 50% ethanol to prevent enzymatic degradation of non-structural carbohydrates (NSC; Chlumska et al., 2022). The remaining material was dried and processed for trait analyses.

We measured 13 key functional traits (sensu Violle et al., 2007), capturing variation in resource use, persistence, and stress tolerance. Aboveground traits included plant height, leaf dry matter content (LDMC), leaf carbon (LCC), nitrogen (LNC), phosphorus (LPC), and stable isotope ratios of carbon (δ^13^C) and nitrogen (δ^15^N), which serve as proxies for water-use efficiency and nitrogen cycling (Farquhar et al., 1989; Robinson, 2001; Liancourt et al., 2020). Belowground traits included root nitrogen and phosphorus content (RNC, RPC) and NSC concentration, representing storage capacity and resource economics. Stable isotope and nutrient concentrations were measured at the UC Davis Stable Isotope Facility using EA-IRMS and spectrophotometry after HClO_4_ digestion (Dolezal et al., 2019b).

NSC concentrations, comprising starch, fructans, and soluble sugars (e.g., glucose, fructose, sucrose, arabitol), were quantified using ethanol extraction, ion-exchange chromatography, and pulsed amperometric detection (Chlumska et al., 2022). Starch was analyzed using the Megazyme total starch assay (Megazyme.com).

To assess anatomical adaptations, we prepared double-stained cross-sections of root collars from a subset of species and individuals. These root collars, located at the root–stem transition, contain annual growth rings that allowed us to determine average vessel size and lignification rate (Doležal et al., 2018; Chondol et al., 2023, 2025). Sections were stained with Astra Blue and Safranin to distinguish parenchyma (storage), lignified tissue (support), and xylem vessels (conduction), following standard protocols (Gärtner & Schweingruber, 2013). High-resolution images were analyzed in ImageJ to quantify the proportional area of each tissue type using 100 randomly placed polygons per quadrant (Doležal et al., 2019a; Binter et al., 2025).

To evaluate ecological strategies across life forms, we assigned each species to one of seven morphological groups based on growth form and clonal architecture: monocarpic herbs, polycarpic forbs with taproots, pleiocorms (non-clonal herbs with deep root crowns), short- and long-rhizomatous clonal plants, compact cushion plants, and woody shrubs (Klimešová et al., 2011). This classification captured key distinctions in stress adaptation and persistence strategies along the elevational and edaphic gradients. Finally, trait data were linked to species’ phylogenetic relationships, reconstructed using a multilocus dataset (ITS, trnT–trnL, matK+trnK, rbcL) and Bayesian inference (MrBayes 3.1.2) based on sequence alignments generated in MAFFT v6.0. The resulting tree was used to compute pairwise phylogenetic distances and explore patterns of functional convergence and divergence across communities.

### 2.3 Data Analyses

To assess trait composition across communities, we calculated community-weighted means (CWMs) for each trait based on species abundances and trait values within each vegetation relevé. Multivariate differences in CWMs among communities were examined using Redundancy Analysis (RDA), with trait data centered and standardized. The significance of the RDA model was evaluated using a Monte Carlo permutation test with 999 iterations.

Additionally, univariate comparisons of CWMs across communities were conducted using one-way ANOVA, with community identity as the explanatory variable for each trait. Functional diversity was quantified for individual traits using the mean pairwise dissimilarity (MPD) weighted by species abundance. To assess whether the observed functional diversity deviated from random expectations, we applied a null model in which species (along with their traits) were randomly assigned to relevés while preserving observed species richness and abundance distributions. This randomization was performed 10,000 times. The full regional species pool comprising all species in the dataset was used for the randomizations, as our study encompassed a single floristic region (Dvorský et al., 2011). From these simulations, we calculated the Standardized Effect Size (SES) for functional diversity as:

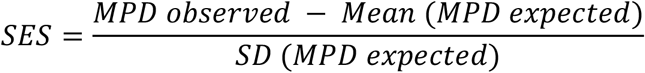

where *MPD observed* is the functional diversity of the observed community, Mean (MPD expected) is the mean of MPD of randomized communities, and SD (MPD expected) is the standard deviation of MPD of randomized communities (Gotelli and McCabe 2002). Differences in SES across communities were first tested using ANOVA, with community identity as a factor. Since SES values varied significantly among communities for all traits (P < 0.05, results not shown), we further tested whether SES values within each community differed significantly from zero using one-sample t-tests. Positive SES values indicate trait divergence (greater trait dissimilarity than expected by chance), negative values indicate trait convergence, and SES values not significantly different from zero suggest neutral assembly patterns. All univariate statistical analyses were performed in R, while multivariate analyses were conducted using Canoco 5 (ter Braak & Šmilauer, 2012).

## 3 Results

### 3.1 Elevational distribution of communities

The elevational distribution of communities reveals a clear environmental gradient shaping community composition and structure (Fig. 2a). Salt marshes (SM), wet grasslands (WE), and shrublands (SH) are predominantly situated at lower elevations (median ∼4500–4700 m a.s.l.), reflecting their preference for relatively less extreme thermal conditions. Steppes (ST) occur slightly higher (median ∼4750 m), overlapping with shrublands but extending into cooler zones. In contrast, screes (SC) and the alpine zone (AL) are restricted to the highest elevations (median ∼5100–5300 m), with the alpine vegetation reaching up to 5850 m a.s.l.—near the upper limits of vascular plant life.

**Figure 2.**
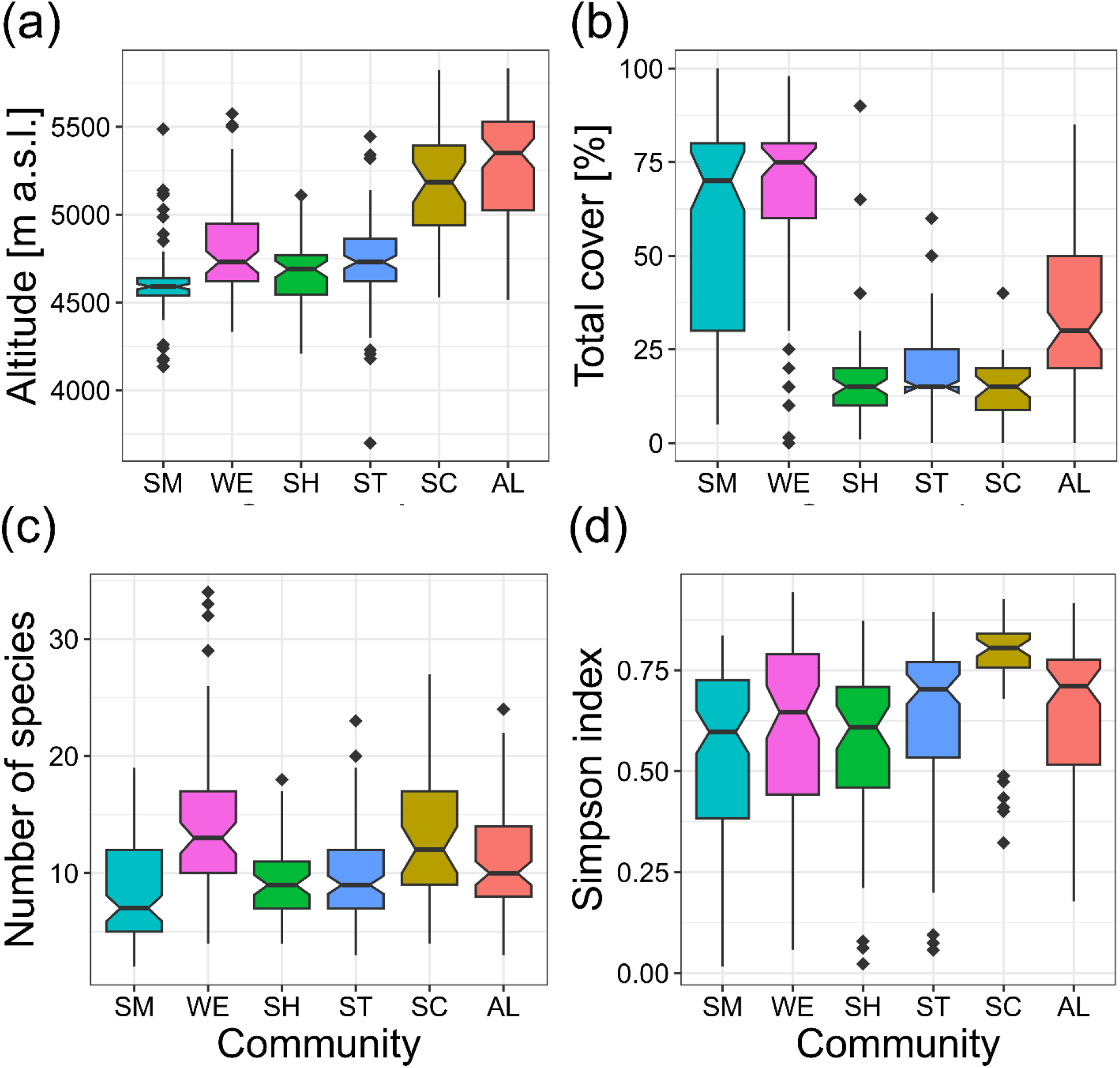
Community characteristics across six Himalayan vegetation types. Boxplots show variation in (a) altitude, (b) total vegetation cover, (c) species richness, and (d) Simpson diversity index across six community types: salt marshes (SM), wet grasslands (WE), shrublands (SH), steppes (ST), screes (SC), and alpine zone (AL). Boxes represent interquartile ranges, horizontal lines show medians, notches indicate 95% confidence intervals, and whiskers extend to 1.5× the interquartile range.

### 3.2 Total vegetation cover across communities

Total vegetation cover varied markedly across the different community types (Fig. 2b), reflecting underlying differences in environmental productivity and disturbance. Wet grasslands (WE) and salt marshes (SM) exhibited the highest total cover (median ∼75%), indicative of high plant productivity in these mesic, nutrient-rich habitats. However, salt marshes showed greater variability, with some plots exhibiting low cover, likely due to spatial heterogeneity in salinity and inundation. Alpine vegetation (AL) also showed moderately high cover (median ∼50%) but with substantial variability, potentially linked to microsite conditions and snow cover persistence. In contrast, shrublands (SH), steppes (ST), and screes (SC) exhibited the lowest vegetation cover (medians ∼20–25%), reflecting a combination of aridity, low nutrient availability, and mechanical instability. Particularly in screes, extreme substrate dynamics and drought likely limit vegetative growth, resulting in sparse plant cover.

### 3.3 Species richness and diversity

Patterns of species richness and diversity further reflect the influence of ecological filters across the studied habitats. Salt marshes (SM) and alpine communities (AL) showed the lowest median species richness (∼10 species per plot, Fig. 2c), consistent with strong environmental filtering under high salinity or cold stress. Wet grasslands (WE) displayed the highest richness (median ∼15 species), with several outliers exceeding 30 species, highlighting their role as biodiversity hotspots under favorable hydrological conditions. Shrublands (SH), steppes (ST), and screes (SC) displayed intermediate levels of species richness, though SC exhibited high variability. In terms of Simpson diversity (Fig. 2d), screes (SC) and the alpine zone (AL) had the highest values (median ∼0.8), suggesting communities dominated by fewer but more evenly abundant species typical of systems shaped by abiotic stress. Salt marshes and shrublands had lower diversity (median ∼0.6–0.65), while wet grasslands and steppes exhibited intermediate diversity (median ∼0.7–0.75), likely due to a mix of stress and biotic interactions. These patterns suggest that although species richness may decline with elevation or disturbance, diversity and evenness can remain high where a few specialized species dominate.

### 3.4 Ordination of trait–community relationships

Redundancy analysis (RDA; F = 5.0, P = 0.002, explained variance = 19.4%) revealed clear functional separation among communities based on their CWM trait profiles (Fig. 3). Salt marshes and wet grasslands grouped along axes associated with clonality, LDMC, LCC, and stable isotope signatures (δ^13^C, δ^15^N), while shrub-dominated steppes and open steppes were aligned with axes reflecting tall stature, lignification, vessel diameter, and nutrient-rich traits such as LNC and NSC. Scree vegetation was distinctly associated with NSC, vessel diameter, and RPC, confirming a unique trait combination linked to extreme edaphic stress. Alpine vegetation was functionally less distinct from the other communities, but shared affinity with conservative traits (e.g., high RPC, low height), suggesting a convergence of trait strategies under cold, short-season conditions.

**Figure 3.**
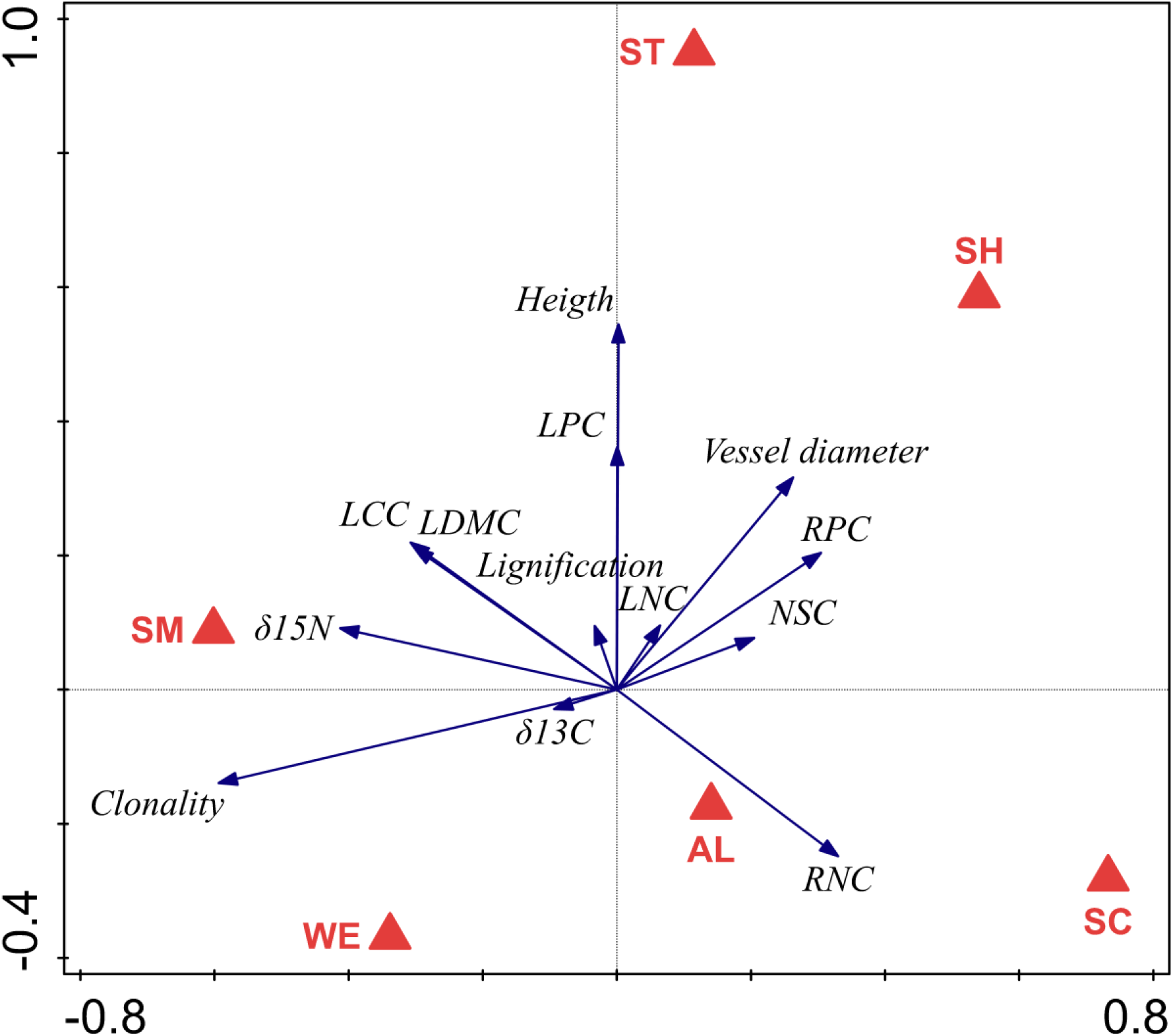
Results of RDA conducted on CWM. (F= 5.0, P = 0.002, explained variability – 19.4%). For abbreviations of communities, see Table 1; for abbreviations of traits, see Table 2.

### 3.5 Community trait differentiation

Community-weighted mean (CWM) values of most measured traits differed significantly among habitat types (Fig. 4; Table 2), except for δ^13^C. Functional difference was most pronounced between halophilous salt marshes and unstable screes (Fig. 2). Salt marshes were dominated by clonal species with high leaf dry matter content (LDMC), leaf carbon concentration (LCC), and enriched δ^15^N signatures, traits associated with stress tolerance under saline and periodically inundated conditions. Wet grasslands, functionally most similar to salt marshes, were characterized by small-statured species with elevated LDMC, LCC, and δ^13^C, consistent with moderate water stress and seasonal saturation.

**Table 1.** Overview of the studied plant communities. Each community is identified by its abbreviation and briefly characterized by dominant vegetation types and habitat conditions.

| Community | Abbreviation | Description |
| --- | --- | --- |
| Salt marshes | SM | Halophilous grasslands |
| Wet grasslands | WE | Grasslands on wetter places, often along the streams |
| Shrublands | SH | Steppe vegetation with high abundance of shrubs |
| Steppes | ST | Steppe vegetation |
| Scree | SC | Vegetation on unstable substrates of scree |
| Alpine zone | AL | High altitude vegetation of alpine zone |

**Table 2:** Summary and description of the studied traits and their difference between the communities tested by ANOVA.

| Trait | Unit | Abbreviation | Function reflected | P communities |
| --- | --- | --- | --- | --- |
| Plant height | cm | Height | Competition-stress trade-off | <10<sup>-3</sup> |
| Leaf dry matter content | mg/g | LDMC | Supporting tissue content | <10<sup>-3</sup> |
| Leaf nitrogen content | % | LNC | Nitrogen limitation | <10<sup>-3</sup> |
| Leaf carbon content | % | LCC | Supporting tissue content | 0.016 |
| Leaf phosphorus content | % | LPC | Phosphorus limitation | <10<sup>-3</sup> |
| Foliar $\delta^{15}\text{N}$ | ‰ | $\delta^{15}\text{N}$ | Nitrogen limitation | <10<sup>-3</sup> |
| Foliar $\delta^{13}\text{C}$ | ‰ | $\delta^{13}\text{C}$ | Water availability stress | 0.512 |
| Root nitrogen content | % | RNC | Nitrogen limitation and storage | <10<sup>-3</sup> |
| Root phosphorus content | % | RPC | Phosphorus limitation and storage | <10<sup>-3</sup> |
| Root non-structural carbohydrates | % | NSC | Carbon storage and resistance to frost | 0.018 |
| Diameter of xylem vessels | μm | Vessels | Trade-off between water use efficiency and cavitation | <10<sup>-3</sup> |
| Degree of xylem lignification | % | Lignification | Supporting tissue content | <10<sup>-3</sup> |
| Ability of clonal growth | yes/no | Clonality | Vegetative reproduction, microsites foraging | <10<sup>-3</sup> |

**Figure 4.**
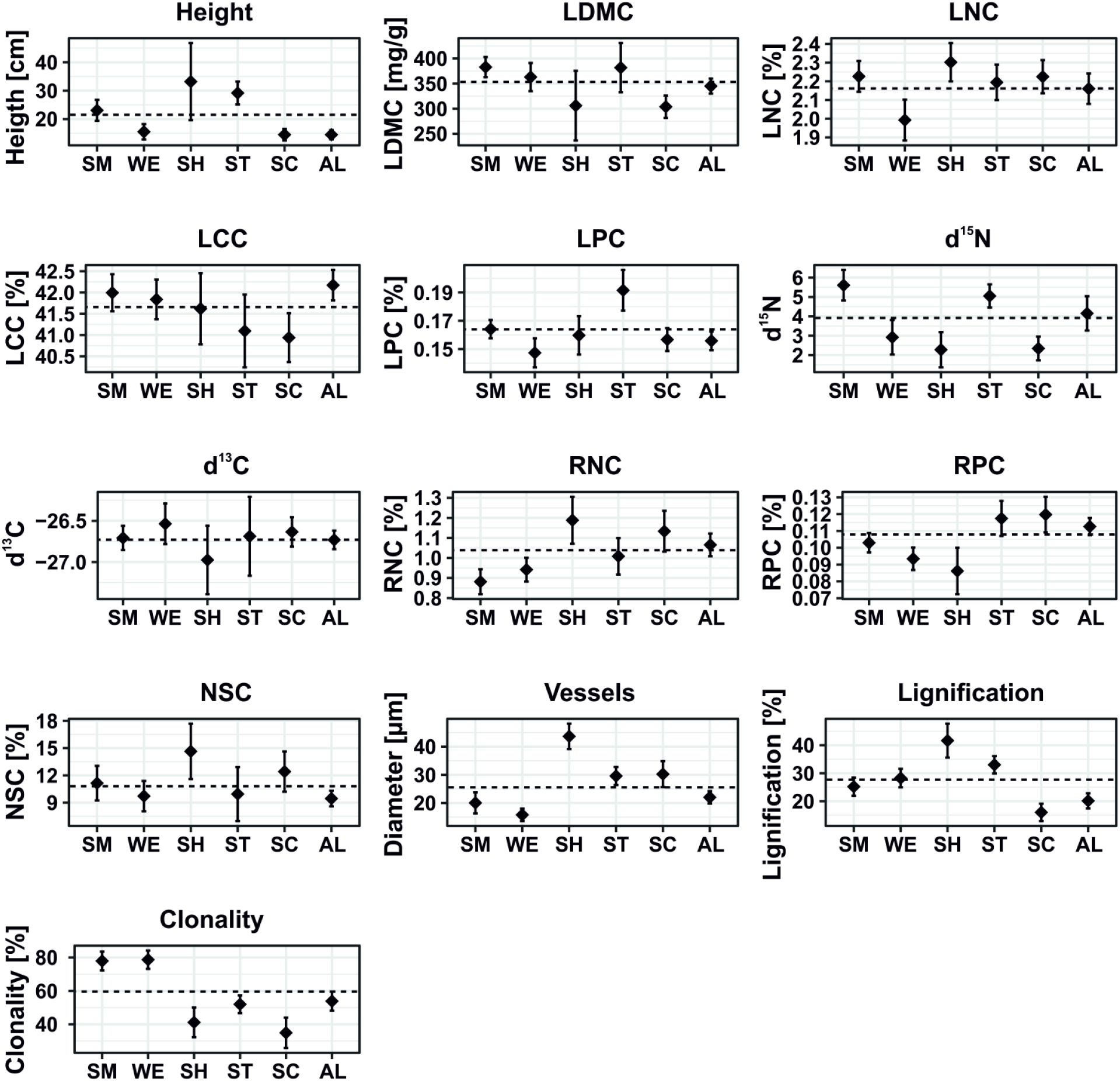
CWM of individual traits in communities. For significant differences, see Table 2. Mean ± 95% confidence interval is displayed. The dashed line shows the average of the trait across all communities. For abbreviations of communities, see Table 1; for abbreviations of traits, see Table 2.

Shrub-dominated steppes and open steppe communities exhibited functional profiles consistent with resource-acquisitive strategies. These habitats were dominated by tall species with high leaf nitrogen concentration (LNC), root nitrogen and phosphorus concentrations (RNC, RPC), non-structural carbohydrates (NSC), and vessel diameter. High levels of lignification were also typical, suggesting structural reinforcement to support taller growth forms and resist desiccation.

In contrast, scree communities, occurring on steep, unstable substrates, were dominated by short-statured species exhibiting high LNC, RNC, RPC, NSC, and large vessel diameters—traits that reflect mechanical resilience, nutrient acquisition efficiency, and belowground carbohydrate storage for survival in physically harsh and nutrient-poor environments. The alpine zone community occupied a central position in trait space, characterized by low plant stature but elevated LCC and RPC, indicating conservative growth strategies and nutrient storage typical of cold, high-elevation environments.

### 3.6 Trait convergence and phylogenetic structure across communities

Standardized effect sizes (SES) revealed distinct patterns of trait convergence and divergence across the six Himalayan plant communities. Overall, SES values revealed that trait convergence was the predominant pattern across most Himalayan communities, while phylogenetic structure varied more widely (Fig. 5). In salt marshes (SM), all traits showed significant convergence, except plant height and δ^15^N, which followed a neutral pattern. Phylogenetic SES, however, indicated significant divergence, suggesting that functionally similar species in this habitat are distantly related, possibly due to strong environmental filtering in saline conditions. In wet grasslands (WE), most traits also exhibited strong convergence, particularly for LDMC, LCC, δ^13^C, and clonality. An exception was LNC, which showed significant divergence, indicating the coexistence of species with different nitrogen-use strategies. Phylogenetic SES was neutral, suggesting limited evolutionary structuring.

**Figure 5.**
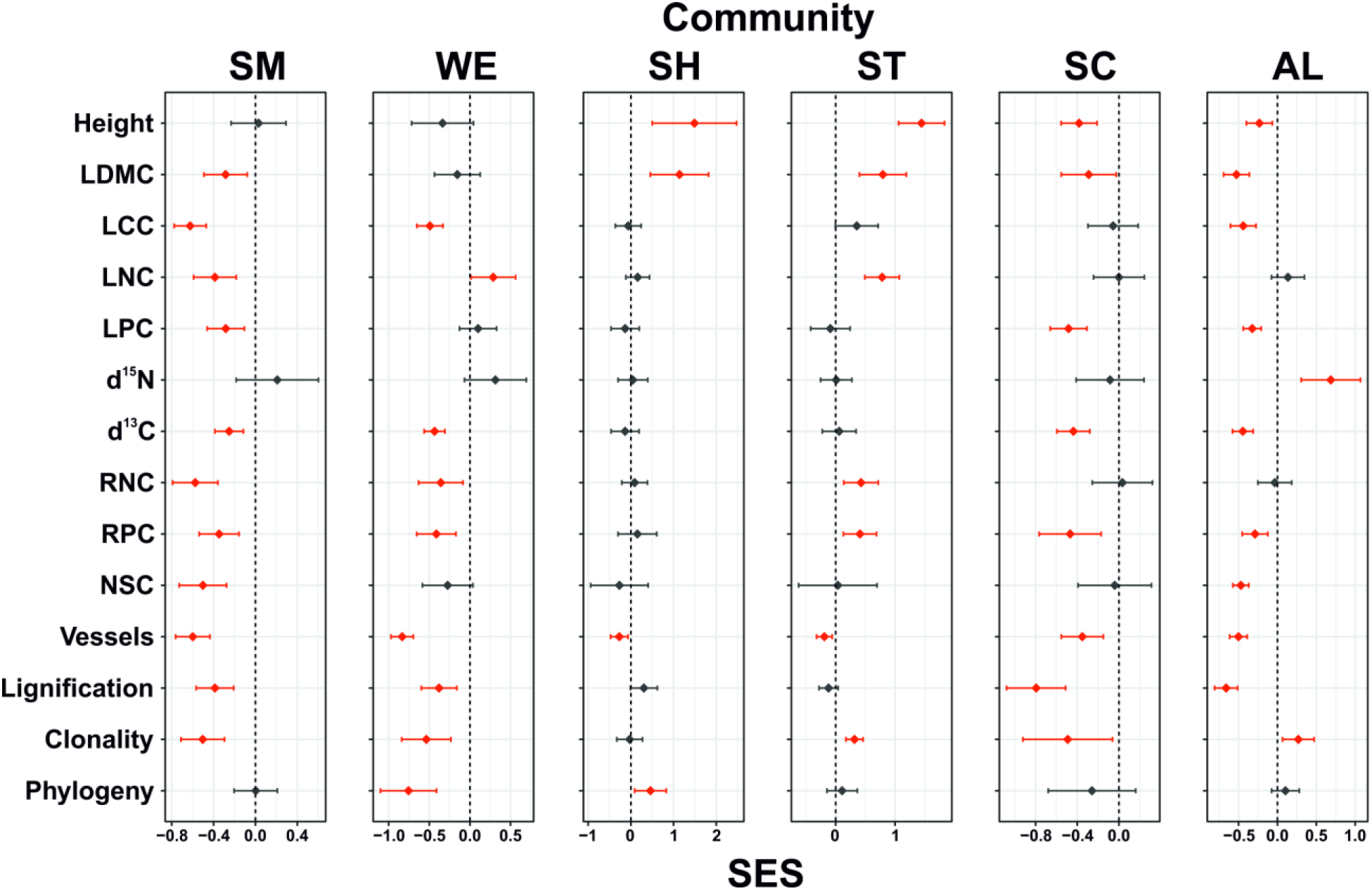
Standardized effect sizes (SES) for individual traits across plant communities. SES values significantly different from zero (P < 0.05, one-sample t-test) are shown in red. Negative SES values indicate trait convergence, positive values indicate trait divergence, and non-significant values (black) reflect neutral patterns. Error bars represent the mean ± 95% confidence interval. The dashed line marks the zero threshold. Community and trait abbreviations are provided in Tables 1 and 2, respectively.

The shrubland (SH) community displayed a more heterogeneous pattern. Most traits showed neutral SES, but plant height and LDMC exhibited significant divergence, while vessel diameter showed convergence. This reflects a community composed of both acquisitive and conservative strategies. Phylogenetic structure was divergent, indicating the coexistence of functionally diverse lineages. Steppes (ST) stood out as the only community where trait divergence predominated. Traits such as height, LDMC, LPC, δ^15^N, RPC, NSC, and lignification all showed positive SES values, indicating a wide range of strategies. Only vessel diameter showed convergence. Phylogenetic SES was significantly divergent, reinforcing the idea of niche partitioning among distantly related taxa in this community.

In screes (SC), all traits with significant SES showed strong convergence, including NSC, LNC, RPC, vessel diameter, and lignification. This trait clustering is likely due to intense abiotic filtering in mechanically unstable, drought-prone habitats. Phylogenetic SES was neutral, implying that environmental pressures selected for similar traits regardless of evolutionary relatedness. The alpine zone (AL) also showed a dominant pattern of trait convergence, particularly for LDMC, LCC, RPC, NSC, and lignification. However, δ^15^N and clonality diverged, suggesting some degree of niche differentiation in nitrogen use and reproductive strategies. Phylogenetic SES indicated divergence, once again highlighting that functionally similar traits have evolved in distantly related species under extreme high-elevation stress. Overall, these results demonstrate that trait convergence predominates in environmentally stressful habitats such as salt marshes, wet grasslands, screes, and alpine zones, while trait divergence emerges under less constrained conditions, as seen in steppes and parts of shrublands. The widespread decoupling between functional and phylogenetic patterns supports the hypothesis of convergent evolution, where unrelated species adopt similar strategies in response to shared environmental filters.

## 4 Discussion

### 4.1 Functional convergence as the dominant assembly pattern

Across the majority of Himalayan communities, functional convergence was the prevailing pattern, particularly for traits linked to stress tolerance and resource conservation such as LDMC, LNC, and RPC. This was most pronounced in environments subject to intense abiotic stress, such as salt marshes, screes, and the alpine zone, where environmental filtering restricts the range of viable ecological strategies. These results are consistent with earlier work showing that strong abiotic filters select a narrow suite of traits in stressful environments (Dvorský et al., 2011;

Spasojevic & Suding, 2012; de Bello et al., 2021). Convergence extended beyond foliar traits to root and stem anatomy and physiology (e.g., lignification, NSC, vessel diameter) in screes and alpine vegetation, indicating a close coupling between anatomical adaptation and habitat harshness (Binter et al., 2025). Comparable convergence in conservative traits, such as high tissue density or low SLA, has been reported from other mountain and stress-prone systems worldwide (Wright et al., 2004; Ratier-Backes et al., 2023).

Evidence from other biomes reinforces this pervasiveness of convergence under abiotic stress. Liu et al. (2013) documented consistent convergence in subtropical forests under hydrological limitation, while Mason et al. (2011) and Lv et al. (2024) highlighted environmental filtering as a driver of low trait dispersion but high phylogenetic diversity. de Bello et al. (2011) showed that Himalayan stress-prone habitats promote narrow trait ranges, particularly in root and clonality traits. In the Apennines, Stanisci et al. (2020) found strong convergence in leaf traits in alpine vegetation exposed to cold and wind, and Scherrer et al. (2019) demonstrated the role of habitat filtering and stature in shaping high-elevation convergence in the Swiss Alps. Mechanistic evidence comes from the anatomical study of Himalayan graminoids by Doležal et al. (2019b), which revealed convergent development of mechanical support and water storage structures in unrelated species across wide elevational and hydrological gradients. A similar pattern was observed in European wetland dicots (Doležal et al., 2021), where phylogenetically distinct lineages occupying hydrologically similar habitats repeatedly evolved analogous anatomical and morphological traits. Together, these studies demonstrate that strong environmental constraints can override phylogenetic signals, repeatedly producing similar morpho-anatomical strategies in unrelated clades.

### 4.2 Divergence in productive habitats

Against this background of widespread convergence, more productive Himalayan communities, such as steppes and shrublands, displayed pronounced divergence in traits linked to stature, LDMC, and nutrient content. In steppes, high divergence in LNC and RPC aligns with studies showing that reduced abiotic stress and intensified biotic interactions promote greater trait dispersion (Götzenberger et al., 2012; Mason et al., 2011). The co-occurrence of acquisitive and conservative strategies in these systems suggests stabilizing niche differentiation, where contrasting resource-use strategies persist through spatial or temporal partitioning. Wedel et al. (2024) similarly found that encroaching shrubs in grasslands adopt divergent strategies, with acquisitive species maximizing rapid growth and conservative species investing in persistence, altogether facilitating coexistence in heterogeneous environments. Comparable divergence is reported in mid-successional temperate and tropical forests, where complementary strategies drive coexistence (Lanta et al., 2023).

The shift from convergence under stress to divergence under higher productivity mirrors findings by Gross et al. (2013), who showed that environmental filtering dominates in arid environments at broad scales, but competitive exclusion in productive microsites fosters local divergence. Mudrák et al. 2015) observed a similar transition in wet meadows, from clustering in low-productivity sites to divergence in high-productivity patches, driven by competitive exclusion. In our dataset, steppes and shrublands followed this pattern, indicating a shift from abiotic filtering to competitive niche differentiation along productivity gradients (Carboni et al., 2014).

### 4.3 Phylogenetic divergence with trait convergence

A striking feature of many Himalayan communities was the decoupling of functional and phylogenetic diversity. Stress-prone habitats, such as salt marshes, screes, alpine zone, exhibited strong convergence in LDMC, LCC, RPC, NSC, and vessel diameter, while phylogenetic SES values were divergent or neutral. Salt marshes showed convergence across most traits yet significant phylogenetic overdispersion, whereas screes were functionally convergent but phylogenetically neutral. This pattern indicates that functionally similar strategies evolved independently in distantly related lineages under parallel environmental selection.

These findings align with the global synthesis by Hähn et al. (2023), which demonstrated that decoupling between functional and phylogenetic diversity is common in communities subject to strong environmental filters, particularly in mountains and drylands. Similar trait–phylogeny decoupling was reported by Choler (2005) in the southwestern Alps, where gradients in snowmelt timing and zoogenic disturbance generated predictable shifts in leaf traits and diversity patterns.

The anatomical convergence we observed in screes and alpine vegetation mirrors that found in Himalayan graminoids (Doležal et al., 2019) and European wetland dicots (Doležal et al., 2021), reinforcing that shared environmental pressures, not common ancestry, shape functional similarity. In subtropical forests, Liu et al. (2013) likewise found trait dispersion more strongly related to environmental and spatial factors than to phylogeny, and He et al. (2022) showed that traits such as SLA and wood density better predicted demographic performance than phylogenetic position. In our steppe communities, however, high divergence coincided with phylogenetic overdispersion, suggesting competitive exclusion and limiting similarity as assembly drivers (Gross et al., 2013; Erdős et al., 2024).

### 4.4 Ecological filters and trait-specific responses across habitats

Trait–environment relationships across Himalayan vegetation types revealed the distinct ecological filters structuring each community. In screes and alpine zones, steep slopes and mechanical instability favoured low-statured species with robust anatomical features and elevated non-structural carbohydrate (NSC) concentrations, enhancing persistence under recurrent physical disturbance. Salt marshes and wet grasslands were dominated by clonal species with high LDMC and LCC, reflecting adaptations to salinity, seasonal waterlogging, and vegetative spread where seedling establishment is constrained (de Bello et al., 2011). Shrublands combined divergence in traits related to stature and leaf tissue investment with convergence in xylem traits, indicating simultaneous filtering by competition and water limitation. These habitat-specific patterns highlight the importance of incorporating anatomical and belowground traits to capture the full spectrum of adaptive responses in heterogeneous environments (Klimešová et al., 2024; Binter et al., 2025). Recent work clarifies the mechanisms underpinning such trait distributions in stressful mountain systems. Chlumská et al. (2023) demonstrated that high-elevation species maintain elevated NSC concentrations as a carbon reserve to buffer severe cold and short growing seasons, while Doležal et al. (2024) showed that biomass allocation strategies among Himalayan plants align with optimal partitioning theory: alpine species invest preferentially in leaves and basal stem structures to maximise carbon gain and storage, whereas steppe species allocate more to deep roots to enhance water uptake.

Clonal growth emerged as a key trait influencing assembly processes across communities. In line with de Bello et al. (2011), we found that clonality was particularly prevalent in low-stature, stress-prone habitats such as salt marshes and alpine zones, suggesting its role in persistence and vegetative spread where seedling establishment is constrained. However, divergence in clonality (e.g., in the alpine zone) also points to niche partitioning among co-occurring clonal types, as previously observed in East Ladakh (de Bello et al. 2011). These findings reinforce the view that clonality is shaped by both abiotic filtering and competitive dynamics, and highlight the need to analyze clonal traits with sufficient resolution to distinguish their multifaceted roles in assembly. Recent work has also emphasized the adaptive value of clonal integration and plasticity in environments with patchy resources or recurrent disturbance (Klimešová & Herben, 2024).

### 4.5 Linking species richness, functional composition, and trait dispersion

Patterns of species richness, trait dispersion (SES), and community-weighted mean (CWM) trait values varied markedly among Himalayan vegetation types, reflecting differences in environmental filtering and competitive interactions. Strongly filtered habitats, such as salt marshes and alpine tundra, were species-poor, showed pronounced trait convergence (negative SES), and often exhibited low CWM values for stature but high values for root nutrient traits (RNC, RPC), indicating stress-adapted, resource-conserving strategies. In contrast, species-rich communities such as wet grasslands and steppes displayed greater trait divergence (positive SES) and higher CWMs for acquisitive traits (e.g. LNC, RPC), consistent with niche partitioning in more productive or less stressful environments.

These findings align with the view that interpreting trait dispersion requires integration with CWMs (de Bello et al., 2016; Götzenberger et al., 2016). In alpine and halophytic habitats, strong abiotic filters selected for compact, low-stature growth forms with limited variation (low SES, low CWM for height). Conversely, in steppes, competition from tall dominants likely increased community mean height (high CWM) while divergence in other traits (positive SES) arose from complementary strategies such as variation in rooting depth or nutrient uptake. This scale-dependent interaction between broad climatic filters (setting dominant CWMs) and fine-scale heterogeneity (modulating functional diversity) mirrors patterns described by de Bello et al. (2013).

Asymmetric competition can also decouple CWMs and SES, elevating CWM trait values while simultaneously reducing dispersion if only one strategy (e.g., tall stature or early flowering) is viable under strong selection. This phenomenon is analogous to experimental results from temperate systems (Doležal et al., 2011), where disturbance (e.g., mulching and mowing) enabled co-occurrence of tall and short grassland plants (high SES, low CWM), while undisturbed abandoned meadows were dominated by uniformly tall species (high CWM, low SES). A similar pattern emerged in Himalayan shrublands, where many traits showed neutral SES with intermediate CWMs, potentially reflecting the competitive dominance of functionally similar but phylogenetically distant taxa—such as *Caragana versicolor, Lonicera spinosa, Krascheninikovia pungens, Artemisia santolinifolia*, and *Scrophularia dentata*—which may exclude alternative strategies and maintain functional homogeneity despite phylogenetic diversity.

Importantly, the same assembly process can yield contrasting patterns across different traits. In alpine and scree communities, conservative traits such as high RPC and NSC converged strongly, whereas clonality and δ^15^N diverged, indicating that environmental filtering and niche differentiation can act simultaneously on different functional dimensions. Comparable patterns were observed in mulched temperate meadows (Doležal et al., 2011), where SLA converged due to nutrient enrichment, but clonality and belowground traits diverged through complementary resource-use strategies. These results reinforce the trait-specific nature of community assembly and the need to examine SES and CWM jointly. High SES may indicate niche complementarity in more productive systems (e.g. steppes), while low SES with low CWMs reflects severe abiotic filtering (e.g., salt marshes). Pooling traits into a single diversity index risks masking such signals, as seen in salt marshes, where convergence in LDMC and NSC coincided with low richness and CWM variance, yet divergence in δ^15^N and clonality revealed persistent fine-scale heterogeneity in nutrient use and reproductive strategies.

## 5 Conclusions and implications for trait-based community ecology in mountain systems

Our study reveals that trait convergence dominates across Himalayan plant communities experiencing strong abiotic filtering, while trait divergence becomes more prominent in more productive or less stressful environments. These patterns indicate that environmental and biotic filters act in parallel and often trait-specific ways. Traits associated with persistence under stress, such as LDMC, NSC, and root phosphorus content, tended to converge in species-poor habitats, whereas traits linked to competition or reproductive strategies, such as clonality or δ^15^N, often diverged, even within the same communities.

By incorporating a broad suite of above- and belowground traits—including stem anatomical traits, clonal growth forms, and non-structural carbohydrate storage—we uncovered key dimensions of ecological strategy that are often overlooked. The frequent decoupling between functional and phylogenetic diversity indicates that unrelated lineages independently converge on similar strategies in response to shared environmental constraints. This highlights the role of evolutionary convergence in shaping plant communities across heterogeneous mountain landscapes. Our findings also demonstrate the utility of combining standardized effect sizes (SES) and community-weighted means (CWMs) to distinguish between assembly processes. While convergence reflects strong environmental filtering and reduced functional variation, divergence often signals niche partitioning and coexistence. Importantly, interpreting these metrics in relation to species richness clarifies whether competitive exclusion, limiting similarity, or stress filtering is the dominant force in a given habitat.

Altogether, this study illustrates the power of integrating multidimensional trait data with phylogenetic frameworks to disentangle the ecological and evolutionary drivers of community assembly. By explicitly linking trait strategies to environmental gradients in one of the world’s most topographically and climatically complex regions, we provide a blueprint for predicting how mountain plant communities may respond to ongoing environmental change.

## Author contributions

JD conceived the ideas and designed the methodology; JD and OM collected the data; OM analysed the data; JD and OM led the writing of the manuscript.

## Acknowledgements

We thank Miroslav Dvorský, Zuzana Chlumská, Pierre Liancourt, Lars Götzenberger, Martin Kopecký, Martin Macek and Klara řeháková for their invaluable support with field sampling and assistance with data analyses.

## Data Availability Statement

Data available from the Zenodo repository doi: XXX (Dolezal & Mudrak, 2025).

## References

1. Baraloto, C., Hardy, O.J., Paine, C.E.T., Dexter, K.G., Cruaud, C., Dunning, L.T., Gonzalez, M.A., Molino, J.F., Molino, D., Savolainen, V., and Chave, J. (2012). Using functional traits and phylogenetic trees to examine the assembly of tropical tree communities. J. Ecol. 100: 690–701. 10.1111/j.1365-2745.2012.01966.x

2. Binter, J., Macek, M., and Doležal, J. (2025). Integrating morphological, anatomical, and physiological traits to explain elevational distributions in Himalayan steppe and alpine plants. J. Integr. Plant Biol. 00: 1–15. 10.1111/jipb.13971

3. Carboni, M., de Bello, F., JaneČek, Š., Doležal, J., Horník, J., Lepš, J., Reitalu, T., and Klimešová, J. (2014). Changes in trait divergence and convergence along a productivity gradient in wet meadows. Agric. Ecosyst. Environ. 182: 96–105. 10.1016/j.agee.2013.12.014

4. Chlumská, Z., Liancourt, P., Hartmann, H., Bartoš, M., Altman, J., Dvorský, M., HubáČek, T., Borovec, J., Čapková, K., Kotilínek, M., and Doležal, J. (2022). Species- and compound-specific dynamics of nonstructural carbohydrates toward the world’s upper distribution of vascular plants. Environ. Exp. Bot. 201: 104985. 10.1016/j.envexpbot.2022.104985

5. Choler, P. (2005). Consistent shifts in alpine plant traits along a meso topographic gradient. Arctic, Antarctic, and Alpine Research 37:444–453.

6. Chondol, T., Klimeš, A., Altman, J., Čapková, K., Dvorský, M., Hiiesalu, I., … and Doležal, J. (2023). Habitat preferences and functional traits drive longevity in Himalayan high-mountain plants. Oikos 2023(10): e010073.

7. Chondol, T., Klimeš, A., Hiiesalu, I., Altman, J., Čapková, K., Jandová, V., Kopecký, M., Macek, M., řeháková, K., and Doležal, J. (2025). Contrasting habitat associations and ecophysiological adaptations drive interspecific growth differences among Himalayan high-mountain plants. Ann. Bot. mcaf014. 10.1093/aob/mcaf014

8. Craven, D., Eisenhauer, N., Pearse, W.D., et al. (2018). Multiple facets of biodiversity drive the diversity–stability relationship. Nat. Ecol. Evol. 2: 1579–1587.

9. de Bello, F., Carmona, C.P., Dias, A.T.C., Götzenberger, L., Moretti, M., and Berg, M.P. (2021). Handbook of trait-based ecology: From theory to R tools. Cambridge University Press. 10.1017/9781108628426

10. de Bello, F., Carmona, C.P., Lepš, J., Szava-Kovats, R., and Pärtel, M. (2016). Functional diversity through the mean trait dissimilarity: resolving shortcomings with existing paradigms and algorithms. Oecologia 180: 933–940. 10.1007/s00442-016-3546-0

11. de Bello, F., Doležal, J., Ricotta, C., Klimešová, J., and Chytrý, M. (2011). Plant clonal traits and species diversity in the subnival zone of the Western Himalayas. Preslia 83: 455–471. 10.23855/preslia.2011.455

12. de Bello, F.D., Lavorel, S., Lavergne, S., Albert, C.H., Boulangeat, I., Mazel, F., and Thuiller, W. (2013). Hierarchical effects of environmental filters on the functional structure of plant communities: a case study in the French Alps. Ecography 36: 393–402. 10.1111/j.1600-0587.2012.07438.x

13. Díaz, S., Lavorel, S., de Bello, F., Quétier, F., Grigulis, K., and Robson, M. (2007). Incorporating plant functional diversity effects in ecosystem service assessments. Proc. Natl. Acad. Sci. USA 104: 20684–20689. 10.1073/pnas.0704716104

14. Doležal, J., Chondol, T., Chlumská, Z., Altman, J., Čapková, K., Dvorský, M., Fibich, P., Korznikov, K.A., Ruka, A.T., Macek, M., and řeháková, K. (2024). Contrasting biomass allocations explain adaptations to cold and drought in the world’s highest-growing angiosperms. Ann. Bot. 134: 401–414. 10.1093/aob/mcae028

15. Doležal, J., Dvorský, M., Börner, A., Wild, J., and Schweingruber, F.H. (2018). Anatomy, Age and Ecology of High Mountain Plants in Ladakh, the Western Himalaya. Springer, Cham.

16. Dolezal, J., Dvorsky, M., Kopecky, M., Altman, J., Mudrak, O., Capkova, K., Rehakova, K., Macek, M., & Liancourt, P. (2019c). Functionally distinct assembly of vascular plants colonizing alpine cushions suggests their vulnerability to climate change. Annals of Botany, 123(4), 569–578. 10.1093/aob/mcy207

17. Doležal, J., Klimeš, A., Dvorský, M., říha, P., Klimešová, J., and Schweingruber, F. (2019a). Disentangling evolutionary, environmental and morphological drivers of plant anatomical adaptations to drought and cold in Himalayan graminoids. Oikos 128: 1576–1587. 10.1111/oik.06451

18. Doležal, J., Kopecký, M., Dvorský, M., Macek, M., řeháková, K., Čapková, K., Borovec, J., Schweingruber, F., Liancourt, P., and Altman, J. (2019b). Sink limitation of plant growth determines treeline in the arid Himalayas. Funct. Ecol. 33: 553–565.

19. Doležal, J., KuČerová, A., Jandová, V., Klimeš, A., říha, P., Adamec, L., and Schweingruber, F.H. (2021). Anatomical adaptations in aquatic and wetland dicot plants: Disentangling the environmental, morphological and evolutionary signals. Environ. Exp. Bot. 187: 104495.

20. Doležal, J., Mašková, Z., Lepš, J., Steinbachová, D., de Bello, F., Klimešová, J., Tackenberg, O., Zemek, F., and Květ, J. (2011). Positive long-term effect of mulching on species and functional trait diversity in a nutrient-poor mountain meadow in Central Europe. Agric. Ecosyst. Environ. 145: 10–28. 10.1016/j.agee.2011.01.010

21. Dvorský, M., Altman, J., Kopecký, M., Chlumská, Z., řeháková, K., Janatková, K., and Doležal, J. (2015). Vascular plants at extreme elevations in eastern Ladakh, northwest Himalayas. Plant Ecol. Divers. 8: 571–584.

22. Dvorský, M., Doležal, J., de Bello, F., Klimešová, J., and Klimeš, L. (2011). Vegetation types of East Ladakh: Species and growth form composition along main environmental gradients. Appl. Veg. Sci. 14: 132–147.

23. Dvorský, M., Macek, M., Kopecký, M., Wild, J., and Doležal, J. (2017). Niche asymmetry of vascular plants increases with elevation. J. Biogeogr. 44: 1418–1425.

24. Erdős, L., Ho, K.V., Bede-Fazekas, Á., Kröel-Dulay, G., Tölgyesi, C., Bátori, Z., et al. (2024). Environmental filtering is the primary driver of community assembly in forest–grassland mosaics: A case study based on CSR strategies. J. Veg. Sci. 35: e13228. 10.1111/jvs.13228

25. Funk, J.L., Larson, J.E., Ames, G.M., Butterfield, B.J., Cavender-Bares, J., Firn, J., Laughlin, D.C., Sutton-Grier, A.E., Williams, L., and Wright, J. (2017). Revisiting the Holy Grail: Using plant functional traits to understand ecological processes. Biol. Rev. 92: 1156–1173. 10.1111/brv.12275

26. Garnier, E., and Navas, M.-L. (2012). A trait-based approach to comparative functional plant ecology: Concepts, methods and applications for agroecology. Agron. Sustain. Dev. 32: 365–399. 10.1007/s13593-011-0036-y

27. Garnier, E., Navas, M.-L., and Grigulis, K. (2015). Plant Functional Diversity: Organism Traits, Community Structure, and Ecosystem Properties. Oxford: Oxford University Press. 10.1093/acprof:oso/9780198757368.001.0001

28. Gärtner, H., and Schweingruber, F.H. (2013). Microscopic preparation techniques for plant stem analysis. Remagen, Germany: Verlag Dr. Kessel.

29. Gotelli, N.J., and McCabe, D.J. (2002). Species co-occurrence: A meta-analysis of J.M. Diamond’s assembly rules model. Ecology 83: 2091–2096. https://doi.org/10.1890/0012-9658(2002)083[2091:SCOAMA]2.0.CO;2

30. Götzenberger, L., Botta-Dukát, Z., Lepš, J., Pärtel, M., Zobel, M., and de Bello, F. (2016). Which randomizations detect convergence and divergence in trait-based community assembly? A test of commonly used null models. J. Veg. Sci. 27: 1275–1287. 10.1111/jvs.12452

31. Götzenberger, L., de Bello, F., Bråthen, K.A., et al. (2012). Ecological assembly rules in plant communities—approaches, patterns and prospects. Biol. Rev. 87: 111–127. 10.1111/j.1469-185X.2011.00187.x

32. Grime, J.P. (2006). Plant Strategies, Vegetation Processes, and Ecosystem Properties, 2nd ed. Chichester: Wiley. 464 pp. ISBN: 978-0-470-85040-4.

33. Gross, N., Börger, L., Soriano-Morales, S.I., Le Bagousse-Pinguet, Y., Quero, J.L., García-Gómez, M., Valencia-Gómez, E., and Maestre, F.T. (2013). Uncovering multiscale effects of aridity and biotic interactions on the functional structure of Mediterranean shrublands. J. Ecol. 101: 637–649. 10.1111/1365-2745.12063

34. Hähn, G.J.A., Damasceno, G., Alvarez-Davila, E., et al. (2025). Global decoupling of functional and phylogenetic diversity in plant communities. Nat. Ecol. Evol. 9: 237–248. 10.1038/s41559-024-02589-0

35. Keddy, P.A. (1992). Assembly and response rules: two goals for predictive community ecology. J. Veg. Sci. 3: 157–164. 10.2307/3235676

36. Klimešová, J., and Herben, T. (2024). Belowground morphology as a clue for plant response to disturbance and productivity in a temperate flora. New Phytol. 242: 61–76. 10.1111/nph.19584

37. Klimešová, J., Doležal, J., Dvorský, M., de Bello, F., and Klimeš, L. (2011). Clonal growth forms in eastern Ladakh, Western Himalayas: classification and habitat preferences. Folia Geobot. 46: 191–217.

38. Kraft, N.J.B., Valencia, R., and Ackerly, D.D. (2008). Functional traits and niche-based tree community assembly in an Amazonian forest. Science 322: 580–582. 10.1126/science.1160662

39. Lavorel, S., and Garnier, E. (2002). Predicting changes in community composition and ecosystem functioning from plant traits: revisiting the Holy Grail. Funct. Ecol. 16: 545–556. 10.1046/j.1365-2435.2002.00664.x

40. Le Bagousse-Pinguet, Y., Liancourt, P., Götzenberger, L., de Bello, F., Altman, J., Brozova, V., et al. (2018). A multi-scale approach reveals random phylogenetic patterns at the edge of vascular plant life. Perspect. Plant Ecol. Evol. Syst. 30: 22–30. 10.1016/j.ppees.2017.10.002

41. Liang, S., Dong, M., Jiang, Y., et al. (2025). Untangling community assembly through functional traits and phylogenetic alpha diversity in subtropical karst forests. Ecol. Evol. 15: e71616. 10.1002/ece3.71616

42. Liu, X., Swenson, N.G., Zhang, J., and Ma, K. (2013). The environment and space, not phylogeny, determine trait dispersion in a subtropical forest. Funct. Ecol. 27: 264–272. 10.1111/1365-2435.12018

43. Luo, Y.-H., Liu, J., Tan, S.-L., Cadotte, M.W., Wang, Y.-H., Xu, K., et al. (2016). Trait-based community assembly along an elevational gradient in subalpine forests: quantifying the roles of environmental factors in inter- and intraspecific variability. PLoS ONE 11: e0155749. 10.1371/journal.pone.0155749

44. Lv, T., Ding, H., Wang, N., Xie, L., Chen, S., Wang, D., and Fang, Y. (2024). The roles of environmental filtering and competitive exclusion in the plant community assembly at Mt. Huangshan are forest-type-dependent. Glob. Ecol. Conserv. 51: e02906. 10.1016/j.gecco.2024.e02906

45. Mason, N.W.H., de Bello, F., Doležal, J., and Lepš, J. (2011). Niche overlap reveals the effects of competition, disturbance and contrasting assembly processes in experimental grassland communities. J. Ecol. 99: 788–796. 10.1111/j.1365-2745.2011.01801.x

46. Mayfield, M.M., and Levine, J.M. (2010). Opposing effects of competitive exclusion on the phylogenetic structure of communities. Ecol. Lett. 13: 1085–1093. 10.1111/j.1461-0248.2010.01509.x

47. McGill, B.J., Enquist, B.J., Weiher, E., and Westoby, M. (2006). Rebuilding community ecology from functional traits. Trends Ecol. Evol. 21: 178–185. 10.1016/j.tree.2006.02.002

48. Mudrák, O., JaneČek, Š., Götzenberger, L., Mason, N.W.H., Horník, J., de Castro, I., Doležal, J., Klimešová, J., and de Bello, F. (2016). Fine-scale coexistence patterns along a productivity gradient in wet meadows: shifts from trait convergence to divergence. Ecography 39: 338–348. 10.1111/ecog.01723

49. Ratier Backes, A., Römermann, C., Alexander, J.M., Arévalo, J.R., Keil, P., Padrón-Mederos, M.A., et al. (2023). Mechanisms behind elevational plant species richness patterns revealed by a trait-based approach. J. Veg. Sci. 34: e13171. 10.1111/jvs.13171

50. Scherrer, D., Mod, H.K., Pottier, J., Schmid, B., and Körner, C. (2019). Disentangling the processes driving plant assemblages in mountain grasslands across spatial scales and environmental gradients. J. Ecol. 107: 265–278. 10.1111/1365-2745.13013

51. Spasojevic, M.J., and Suding, K.N. (2012). Inferring community assembly mechanisms from functional diversity patterns: the importance of multiple assembly processes. J. Ecol. 100: 652–661. 10.1111/j.1365-2745.2011.01945.x

52. Stanisci, A., Bricca, A., Calabrese, V., Cutini, M., Pauli, H., Steinbauer, K., and Carranza, M.L. (2020). Functional composition and diversity of leaf traits in subalpine versus alpine vegetation in the Apennines. AoB PLANTS 12: plaa004. 10.1093/aobpla/plaa004

53. Violle, C., Navas, M.-L., Vile, D., Kazakou, E., Fortunel, C., Hummel, I., and Garnier, E. (2007). Let the concept of trait be functional! Oikos 116: 882–892. 10.1111/j.0030-1299.2007.15559.x

54. Wang, J., Wang, Y., Qu, M., Feng, Y., Wu, B., Lu, Q., He, N., and Li, J. (2022). Testing the functional and phylogenetic assembly of plant communities in Gobi deserts of Northern Qinghai–Tibet Plateau. Front. Plant Sci. 13: 952074. 10.3389/fpls.2022.952074

55. Webb, C.O., Ackerly, D.D., McPeek, M.A., and Donoghue, M.J. (2002). Phylogenies and community ecology. Annu. Rev. Ecol. Syst. 33: 475–505. 10.1146/annurev.ecolsys.33.010802.150448

56. Wedel, E.R., Ratajczak, Z., Tooley, E.G., and Nippert, J.B. (2025). Divergent resource-use strategies of encroaching shrubs: Can traits predict encroachment success in tallgrass prairie? J. Ecol. 113: 339–352. 10.1111/1365-2745.14456

57. Wright, I.J., Reich, P.B., Westoby, M., et al. (2004). The worldwide leaf economics spectrum. Nature 428: 821–827. 10.1038/nature024

